# The generation of HepG2 transmitochondrial cybrids to reveal the role of mitochondrial genotype in idiosyncratic drug-induced liver injury: a translational *in vitro* study

**DOI:** 10.1101/2022.03.21.485109

**Authors:** Amy L. Ball, Carol E. Jolly, Jonathan J. Lyon, Ana Alfirevic, Amy E. Chadwick

## Abstract

**Background:** Evidence supports an important link between mitochondrial DNA (mtDNA) variation and adverse drug reactions such as idiosyncratic drug-induced liver injury (iDILI). Here we describe the generation of HepG2-derived transmitochondrial cybrids in order to investigate the impact of mtDNA variation upon mitochondrial (dys)function and susceptibility to iDILI against a constant nuclear background. In this study, cybrids were created to contain mitochondrial genotypes of haplogroup H and haplogroup J for comparison.

**Methods:** Briefly, HepG2 cells were depleted of mtDNA to make rho zero cells before the introduction of known mitochondrial genotypes using platelets from healthy volunteers (n=10), thus generating 10 distinct transmitochondrial cybrid cell lines. The mitochondrial function of each was assessed at basal state and following treatment with compounds associated with iDILI; flutamide, 2-hydroxyflutamide and tolcapone, by ATP assays and extracellular flux analysis.

**Findings:** Overall, baseline mitochondrial function was similar between haplogroups H and J. However, haplogroup specific responses to mitotoxic drugs were observed; haplogroup J was more susceptible to the inhibition of respiratory complexes I and II, and also to the effects of tolcapone, an uncoupler of mitochondrial respiration.

**Conclusions:** This study demonstrates that HepG2 transmitochondrial cybrids can be created to contain the mitochondrial genotype of any individual of interest, thus providing a practical and reproducible system to investigate the cellular consequences of variation in mitochondrial genome against a constant nuclear background. Additionally the results support that that inter-individual variation in mitochondrial genotype and haplogroup may be a factor in determining sensitivity to mitochondrial toxicants.

**Funding:** This work was supported by the Centre for Drug Safety Science supported by the Medical Research Council, United Kingdom (Grant Number G0700654); and GlaxoSmithKline as part of an MRC-CASE studentship (grant number MR/L006758/1).

## Introduction

Drug-induced liver injury (DILI) is a leading cause of acute liver failure in the western world (Bernal and Wendon, 2013; Tujios and Lee, 2018). Although it is rare (19.1 cases per 100 000 inhabitants), it is a major cause of drug withdrawal due to its associated morbidity and mortality (Leise et al., 2014). Drug-induced liver injury can be broadly divided into two categories, idiosyncratic and intrinsic injury. Idiosyncratic DILI (iDILI) is characterised by a complex dose-response relationship, lack of predictivity from the primary pharmacology of a drug and significant interindividual variability. This means that despite being less common than intrinsic injury, iDILI can be viewed as far more costly (Fontana, 2014).

Drug-induced mitochondrial dysfunction is one of the mechanisms implicated in the onset of DILI, supported by the fact that 50% of drugs with a black box warning for hepatotoxicity also contain a mitochondrial liability, with many of these drugs associated specifically with iDILI (Dykens and Will, 2007). It has been hypothesised that mitochondria is an important source of interindividual variation, underpinning susceptibility to iDILI. Specifically, mitochondria contain multiple copies of their own genome, mitochondrial DNA (mtDNA). Furthermore, single nucleotide polymorphisms (SNPs) in mtDNA are often inherited together forming mitochondrial haplogroups. Not only have associations between specific haplogroups and mitochondrial function been determined, but haplogroups have also been associated with specific adverse drug reactions and drug efficacy (Jones et al., 2021a).

The HepG2 cell line is one of the most commonly used preclinical cell lines for the *in vitro* assessment of DILI. However, this homogenous cell line does not encompass interindividual variation, including that of the mitochondrial genome. The importance of assessing the role of interindividual variation on patient susceptibility to adverse drug reactions is well recognised and the identification of mtDNA variants that confer susceptibility to iDILI could prove invaluable in drug development and drug safety. However, given the high economic cost associated with large, multicentre clinical trials, there is a need for new strategies enabling interindividual variation to be accounted for at the preclinical stage (Fermini et al., 2018; Jones et al., 2021b).

The generation of transmitochondrial cybrids enables mtDNA variation to be incorporated into reproducible *in vitro* models. Transmitochondrial cybrids are typically produced by the fusion of enucleated cells (cytoplasts) or anucleate cells (e.g. platelets) with cells that have been depleted of their mtDNA (rho zero [ρ0] cells) (King and Attardi, 1989). When cybrids are generated using the same population of ρ0 cells fused with anucleate cells harbouring different mtDNA variants, it is possible for the effects of mtDNA to be assessed against a stable nuclear background (Wilkins et al., 2014; Penman et al., 2020). Excitingly, the representation of mtDNA variation using HepG2 transmitochondrial cybrids may offer enhanced preclinical prediction of iDILI by facilitating the *in vitro* elucidation of the mechanistic basis of any differences associated with mtDNA variants, yet to-date there have been no reports of the generation of cybrids from a HepG2 p0 cell line (Bale et al., 2014).

Therefore, the overall aim of this study was to generate a panel of HepG2 transmitochondrial cybrids, as a proof of principle study, to investigate the effect of mtDNA variants upon mitochondrial function and their role, if any, in conferring susceptibility to drug-induced mitochondrial dysfunction. Specifically, ten populations of HepG2 transmitochondrial cybrids (herein referred to as cybrids) were created and characterised; five using platelets derived from healthy volunteers of haplogroup H and five from haplogroup J. Haplogroup H was selected as it is the most common haplogroup in the UK (Eupedia, 2016). Haplogroup J, on the other hand, is less common but is characterised by non-synonymous mutations in regions of the mitochondrial genome that encode mitochondrial respiratory complex I (Van Oven and Kayser, 2009; Eupedia, 2016). See Figure 1 for a schematic overview of the study.

**Figure 1.**
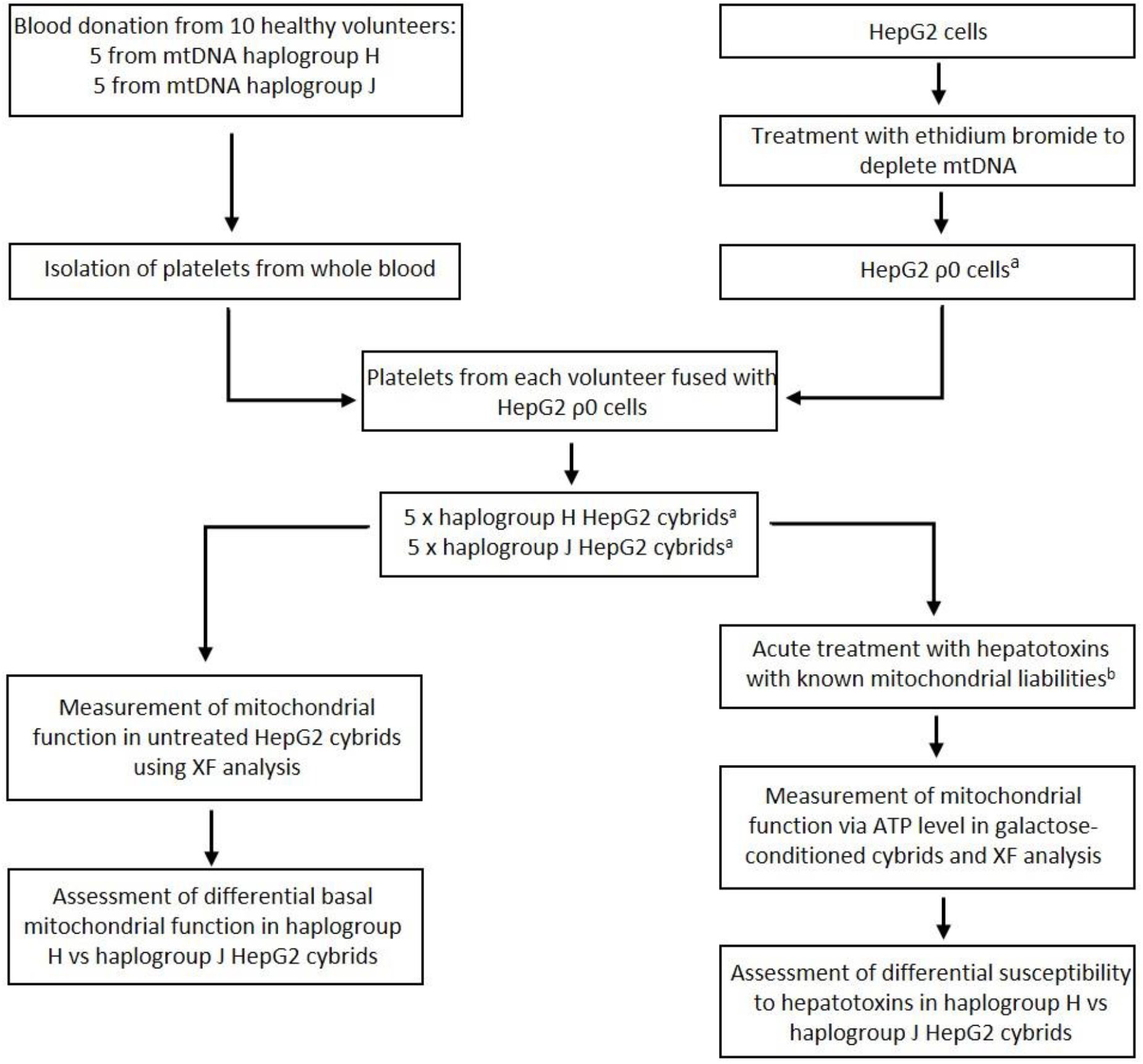
Study Overview. ^a^ HepG2 ρ0 cells were characterised to ensure the complete depletion of mtDNA and HepG2 cybrids were characterised to ensure the expression of mtDNA-encoded proteins. Methods of characterisation are described in the Supplementary Information. ^b^ Flutamide, 2-hydroxyflutamide and tolcapone, alongside non-hepatotoxic structural counterparts; bicalutamide and entacapone. Abbreviations: Abbreviations: mtDNA, mitochondrial DNA; ρ0, rho zero; XF, extracellular flux.

Mitochondrial function was measured by ATP quantification in acutely galactose-conditioned cells, and extracellular flux (XF) analysis of both total respiratory chain function and the function of specific respiratory complexes, in whole and permeabilised cybrids, respectively. Drug-induced mitochondrial dysfunction was assessed by the treatment of cybrids with a panel of compounds selected to comprise of known hepatotoxins (a clinical association with DILI) with proven mitochondrial liabilities; flutamide, 2-hydroxyflutamide and tolcapone, alongside structural counterparts which are not hepatotoxic; bicalutamide and entacapone. Flutamide, a non-steroidal antiandrogen for the treatment of prostate cancer has a boxed warning for hepatotoxicity and is a known inhibitor of mitochondrial complex I (Coe et al., 2007). 2-hydroxyflutamide, is the primary metabolite of flutamide and is a known inhibitor of both respiratory complex I and respiratory complex II. In humans, the rapid first-pass metabolism of flutamide results in 2-hydroxyflutamide having a much higher maximum serum concentration than its parent compound (4400 nM versus 72.2 nM). However, HepG2 cells have limited expression of many enzymes required for xenobiotic metabolism, including cytochrome P450 1A2, the primary route of flutamide metabolism to generate 2-hydroxyflutamide; therefore cybrids were also dosed with 2-hydroxyflutamide directly (Shet et al., 1997; Sison-Young et al., 2015; Ball et al., 2016). Tolcapone, a catechol-o-methyl transferase inhibitor used to treat Parkinson’s disease was withdrawn due to cases of liver failure and is a known uncoupler of oxidative phosphorylation (Olanow, 2000; Benabou and Waters, 2003; Olanow and Watkins, 2007).

## Results

### Haplogroup J cybrids have greater respiratory complex activity than haplogroup H cybrids

Assessment of basal mitochondrial function using a mitochondrial stress test showed no significant difference between haplogroup H and J cybrids in multiple parameters of mitochondrial respiration (Figure 2A). Contrastingly, haplogroup J cybrids had significantly greater (≥2 fold; p=0.025, p=0.001) complex I and II-driven maximal respiration (herein referred to as complex activity for clarity) (Figure 2B).

**Figure 2.**
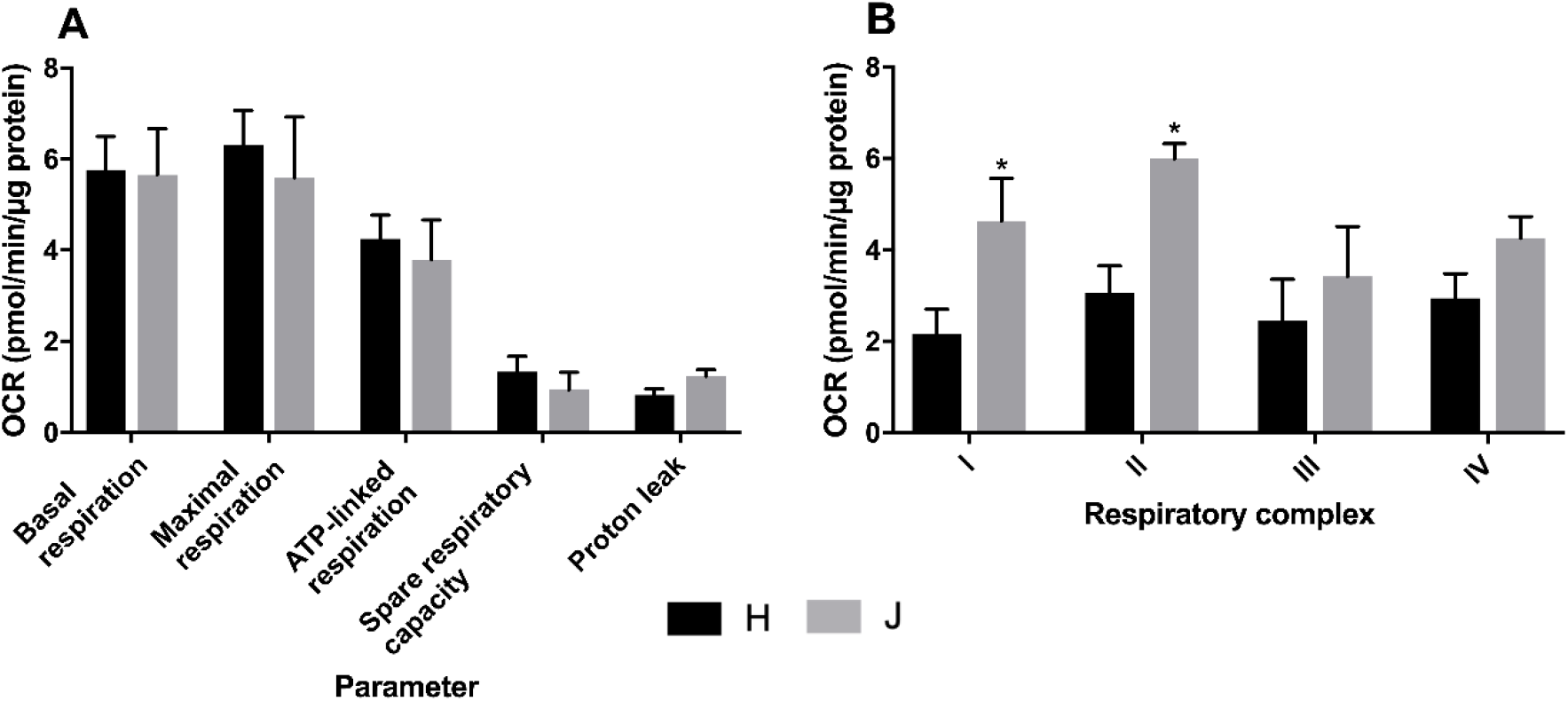
Basal mitochondrial function and respiratory complex activity in haplogroup H and J HepG2 cybrids. **A:** Untreated haplogroup H and J cybrids were assessed using extracellular flux analysis and a mitochondrial stress test to measure: basal respiration, maximal respiration, ATP-linked respiration, spare respiratory capacity and proton leak. **B:** Untreated haplogroup H and J cybrids were assessed using extracellular flux analysis and respiration was stimulated by the supply of respiratory complex-specific substrates. Complex I-IV activity was defined as complex I-IV driven maximal respiration. Statistical significance between haplogroup H and J cybrids: * *p*<0.05. Data are presented as mean + SEM of n=5 experiments. Abbreviations: OCR, oxygen consumption rate. Source data: fig2 – source data file.xslx

### Inhibition of mitochondrial respiration by flutamide is similar in haplogroup H and J cybrids

Acute treatment of galactose-conditioned cybrids with flutamide did not induce cytotoxicity (i.e. no significant lactate dehydrogenase [LDH] release) at any of the concentrations used (data not shown). A substantial, concentration-dependent decrease in ATP was evident when cybrids were treated with ≥33 μM flutamide, but the decrease was not significantly different between haplogroup H and J cybrids (Figure 3A). Concordantly, there was no significant difference in the ATP level EC_50_ between the two haplogroups. XF analysis also revealed no significant difference in parameters of mitochondrial respiration between haplogroup H and haplogroup J cybrids when treated with flutamide, though haplogroup J cybrids did have a greater proton leak and reduced spare respiratory capacity than haplogroup H at all concentrations, but this was not significant (Figure 3B-F).

**Figure 3.**
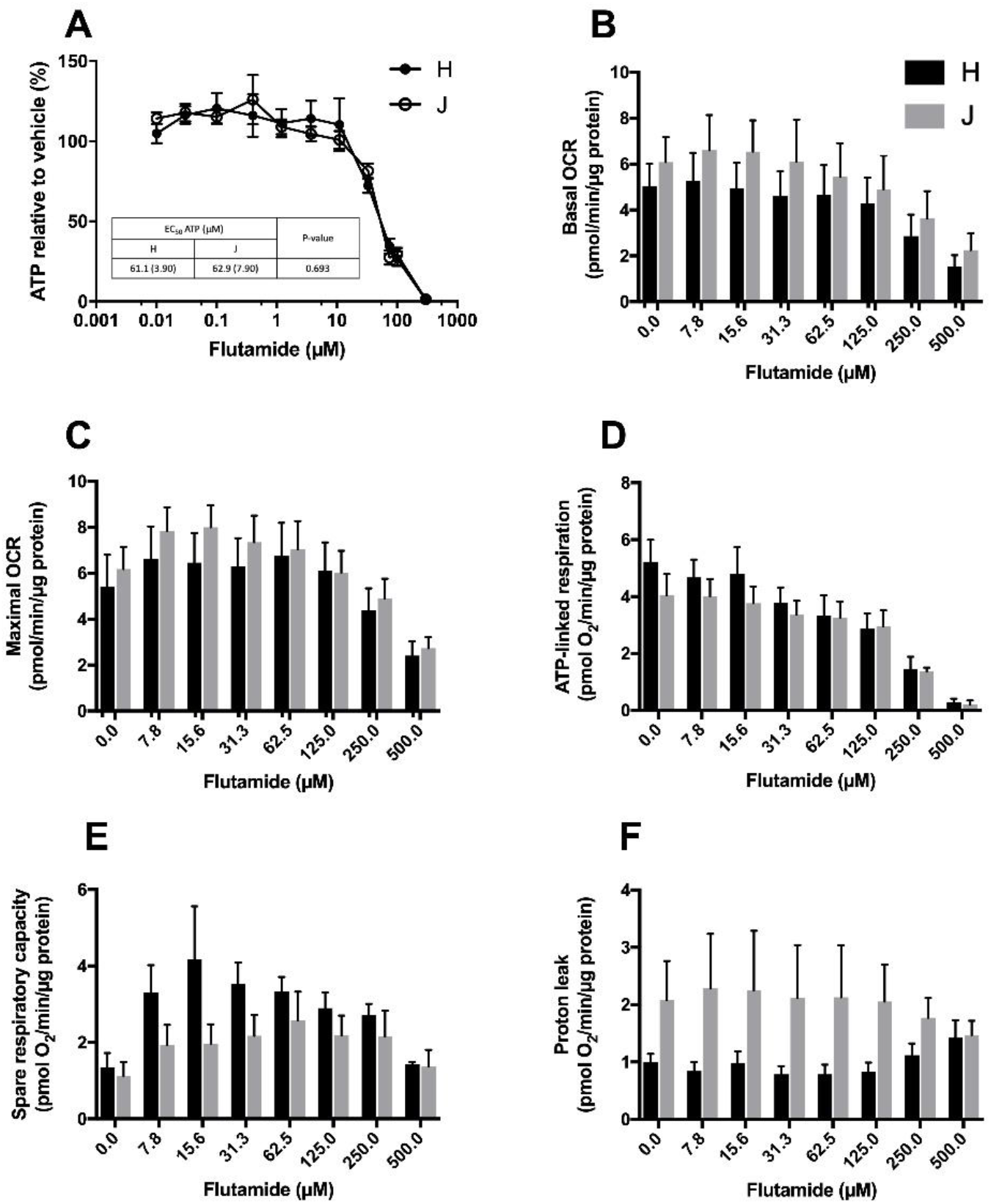
The effect of flutamide upon ATP levels and mitochondrial respiratory function in haplogroup H and J HepG2 cybrids. **A:** Cybrids were treated (2 h) with up to 300 μM flutamide in galactose medium. ATP values are expressed as a percentage of those of the vehicle control. **B-F:** Changes in basal respiration, maximal respiration, ATP-linked respiration, spare respiratory capacity and proton leak following acute treatment with flutamide (up to 500 μM). Data are presented as mean ± SEM of n=5 experiments. Abbreviations: OCR, oxygen consumption rate. Source data: fig3 – source data file.xslx

### Inhibition of mitochondrial respiration by 2-hydroxyflutamide is similar in haplogroup H and J cybrids

Acute treatment of galactose-conditioned cybrids with 2-hydroxyflutamide did not induce cytotoxicity at any of the concentrations used (data not shown). A substantial, concentration–dependent decrease in ATP level was evident when cybrids were treated with ≥33 μM 2-hydroxyflutamide, but the decrease was not significantly different between haplogroup H and J cybrids at any single concentration (Figure 4A); however, a comparison of EC_50_ values showed a small but significantly lower EC_50_ value in haplogroup H cybrids (Figure 4A; p=0.003). XF analysis also revealed no significant difference in parameters of mitochondrial respiration between haplogroup H and haplogroup J cybrids when treated with 2-hydroxyflutamide (Figure 4B-F). As was the case with flutamide, 2-hydroxyflutamide-treated haplogroup J cybrids had a greater proton leak and reduced spare respiratory capacity compared with haplogroup H at most concentrations, but this was not significant (Figure 4E, F).

**Figure 4.**
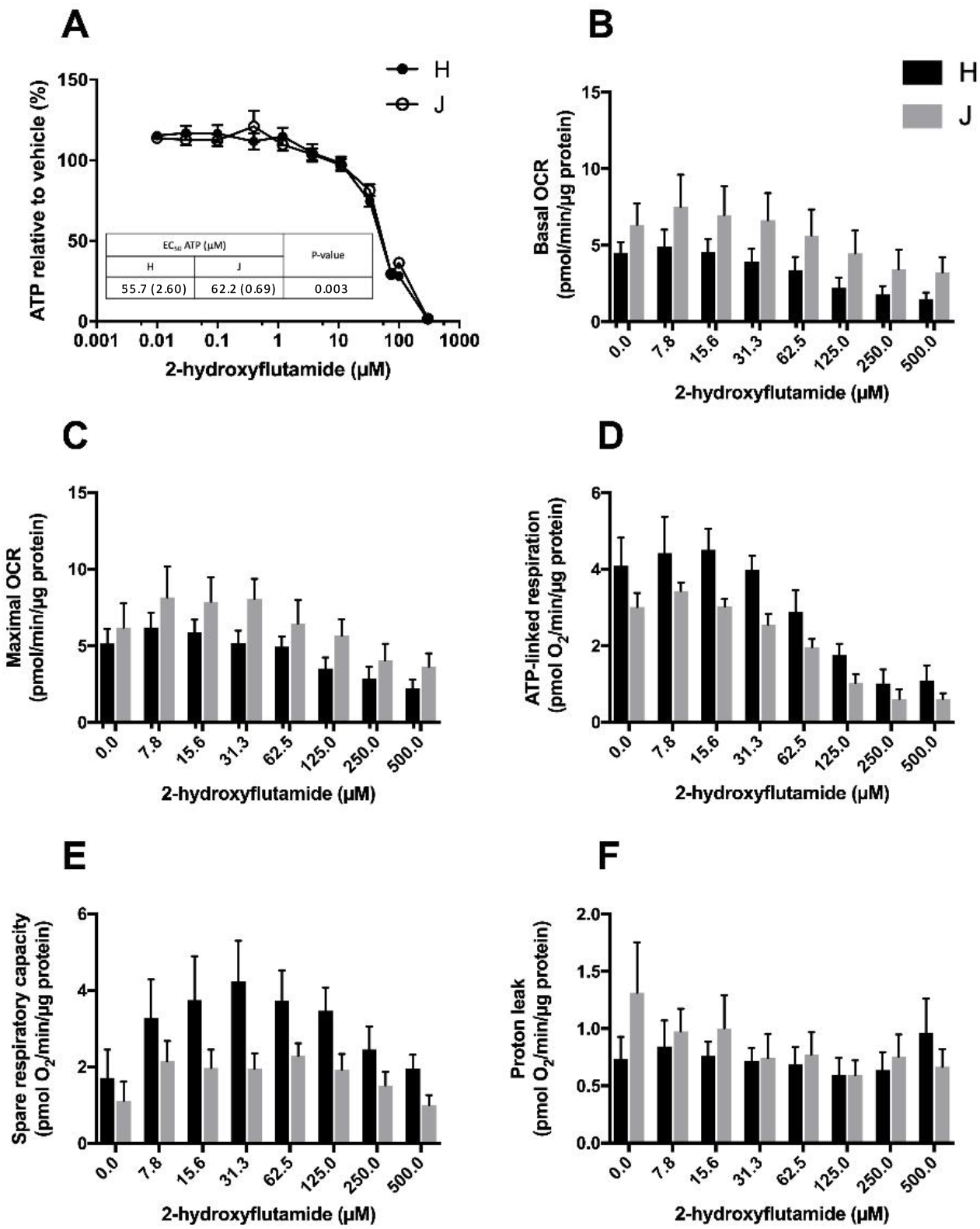
The effect of 2-hydroxyflutamide upon ATP levels and mitochondrial respiratory function in haplogroup H and J HepG2 cybrids. **A:** Cybrids were treated (2 h) with up to 300 μM 2-hydroxyflutamide in galactose medium. ATP values are expressed as a percentage of those of the vehicle control. **B-F:** Changes in basal respiration, maximal respiration, ATP-linked respiration, spare respiratory capacity and proton leak following acute treatment with 2-hydroxyflutamide (up to 500 μM). Data are presented as mean ± SEM of n=5 experiments. Abbreviations: OCR, oxygen consumption rate. Source data: fig4 – source data file.xslx

### Haplogroup H cybrids are resistant to tolcapone-induced ATP depletion at <75 μM

Cellular ATP levels following treatment with up to 75 μM tolcapone were significantly higher in haplogroup H cybrids; this was reflected by the significantly higher EC_50_ in haplogroup H cybrids, almost twice that of haplogroup J (Figure 5A; p<0.0001). At higher concentrations of tolcapone however, there was no significant difference between the two haplogroups (Figure 5A). XF analysis showed no significant difference in parameters of mitochondrial respiration between haplogroup H and haplogroup J cybrids when treated with tolcapone (Figure 5B-F).

**Figure 5.**
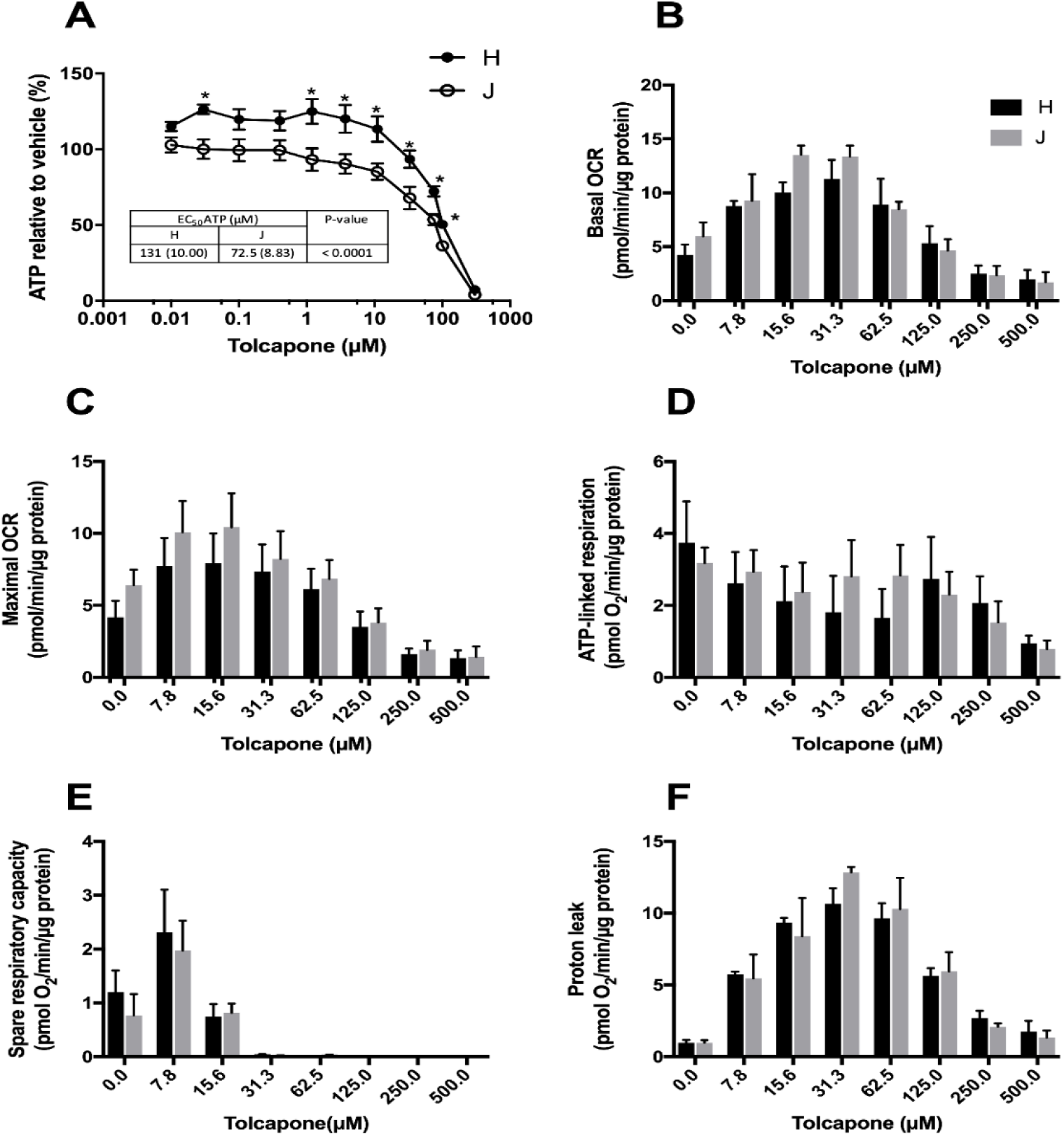
The effect of tolcapone upon ATP levels and mitochondrial respiratory function in haplogroup H and J HepG2 cybrids. **A:** Cybrids were treated (2 h) with up to 300 μM tolcapone in galactose medium. ATP values are expressed as a percentage of those of the vehicle control. **B-F:** Changes in basal respiration, maximal respiration, ATP-linked respiration, spare respiratory capacity and proton leak following acute treatment with tolcapone (up to 500 μM). Statistical significance between: haplogroup H and J cybrids* *p*<0.05. Data are presented as mean ± SEM of n=5 experiments. Abbreviations: OCR, oxygen consumption rate. Source data: fig 5 – source data file.xslx

### Haplogroup J cybrids have greater respiratory complex activity but are more susceptible to inhibition of complex I and II activity by flutamide and 2-hydroxyflutamide

When treated with flutamide (complex I inhibitor) haplogroup J cybrids had significantly greater respiratory complex I/II activity at all but the highest flutamide concentration (250 μM), peaking at approximately two-fold greater activity than haplogroup H cybrids (Figure 6A, B). Though complex I activity was greater in haplogroup J cybrids at all concentrations, haplogroup J cybrids exhibited a bigger decrease (compared with haplogroup H cybrids) in complex I activity relative to control when treated with flutamide (Figure 6A). The effect of treatment with 2-hydroxyflutamide mirrored the effect of its parent compound, flutamide, with greater complex I activity in haplogroup J cybrids at all concentrations, but with haplogroup J cybrids exhibiting the biggest decrease in complex I activity relative to control (Figure 6C).

**Figure 6.**
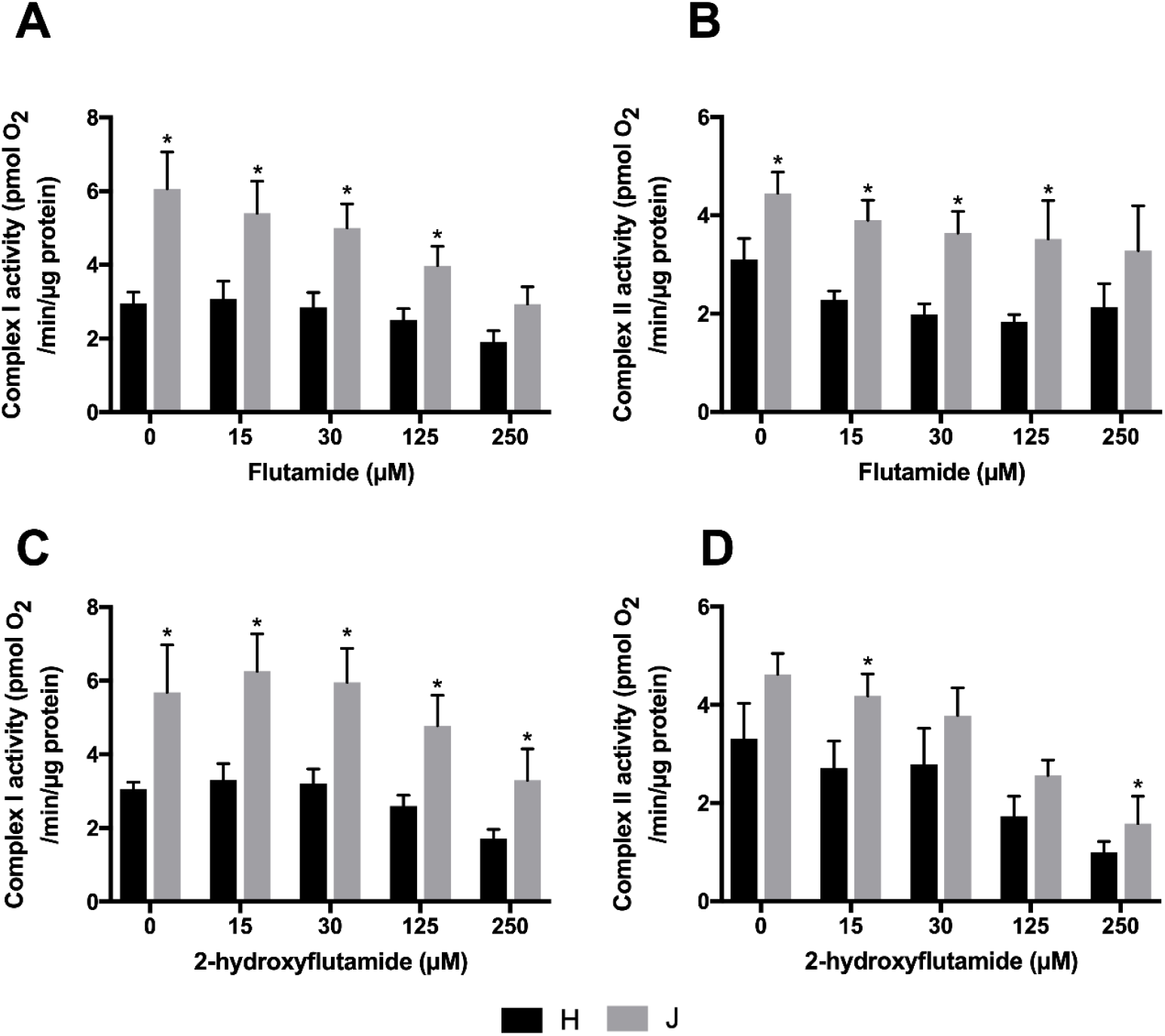
The effect of flutamide and 2-hydroxyflutamide upon respiratory complex I and II in haplogroup H and J HepG2 cybrids. Permeabilised cybrids were acutely treated with flutamide (**A, B**) or 2-hydroxyflutamide (**C, D**) (≤250 μM) before a mitochondrial stress test using extracellular flux analysis. Complex I/II activity was defined as complex I/II driven maximal respiration. Statistical significance between haplogroup H and J cybrids: * p<0.05. Data are presented as mean + SEM of n=5 experiments. Source data: fig 6 – source data file.xslx

A smaller difference was observed in complex II activity between the two cybrid haplogroups when treated with 2-hydroxyflutamide, though greater activity was still evident in haplogroup J cybrids at all concentrations (Figure 6D).

### The impact of bicalutamide and entacapone on mitochondrial respiration and ATP levels is similar in haplogroup H and J cybrids

When treated with bicalutamide and entacapone (non-hepatotoxic structural counterparts of flutamide and tolcapone, respectively), the effect on mitochondrial respiration and ATP levels was similar in haplogroup H and J cybrids. Full results are described in the Supplementary Information.

## Discussion

It has been widely hypothesised that individual variation in mtDNA may be a factor underlying the onset of idiosyncratic adverse drug reactions by drugs known to contain mitochondrial liabilities. Evidence from clinical studies has demonstrated that such associations between mitochondrial genotype and drug efficacy or adverse events do exist (Jones et al., 2021a). However, there is a lack of knowledge and understanding of the importance of these findings due to the presence of nuclear heterogeneity alongside environmental factors in clinical cohorts and the small size of test cohorts, as well as historical limitations in sequencing technology. Because of this we have created a transmitochondrial cybrid cell panel, in which mitochondria of known genotype, sourced from volunteer platelets, were inserted into HepG2 ρ0 cells in order to produce a “personalised model” model of mitotoxicity suitable for mechanistic investigations.

Here we describe the successful generation of 10 distinct transmitochondrial cybrid cell lines, derived from five haplogroup H volunteers and five haplogroup J volunteers. To the best of our knowledge, this is the first time that the creation of a panel of HepG2-derived transmitochondrial cybrids has been reported. Importantly, the HepG2 cell line is one of the most commonly used cell lines in preclinical drug safety testing and therefore the HepG2 cybrids generated in the present study are of great value in improving the understanding of iDILI by providing an *in vitro* representation of the interindividual variation that underpins this adverse drug reaction (Bale et al., 2014). In this work, the utility of these cells to investigate interhaplogroup differences in mitochondrial function has been demonstrated, with HepG2 cybrids of haplogroup J displaying greater mitochondrial respiratory complex activity at basal state. Moreover, our investigations revealed that there were haplogroup-specific differences in susceptibility to hepatotoxic compounds that target the electron transport chain (ETC). Specifically, haplogroup J cybrids were more susceptible to: a reduction in respiratory complex I activity induced by flutamide, a reduction in respiratory complex I and II activity induced by 2-hydroxyflutamide, and a reduction in ATP levels induced by tolcapone.

The methodology developed for the generation of HepG2 cybrids has enabled the creation of a HepG2 cybrid cell panel from two of the most common mitochondrial haplogroups in England, H and J; which account for more than 50% of the population (Eupedia, 2016). Although baseline assessments revealed only marginal differences in mitochondrial function between haplogroup H and J cybrids, the recruitment of healthy volunteers for this study (i.e. no clinical phenotype of mitochondrial dysfunction), meant that the absence of any substantial differences in mitochondrial function was expected. This recruitment of healthy volunteers is representative of the clinical situation, as individuals who experience iDILI tend not to display a phenotype of mitochondrial dysfunction prior to treatment. Additionally, it should be noted that these results were analysed on the basis of macro-haplogroup (i.e H, J). However, each cybrid cell line has been identified as a separate sub-haplogroup (e.g. H1, J2), based upon the accumulation of specific SNPs (see Supplementary Information). This lack of homogeneity within the test groups may mask associations.

The cybrid cell panel was interrogated using known drug mitotoxicants which elicit different effects on the ETC; flutamide and 2-hydroxyflutamide, direct ETC inhibitors, and tolcapone, an ETC uncoupler. The impact of flutamide (complex I inhibitor) and 2-hydroxyflutamide (complex I and II inhibitor) upon cellular ATP level was similar between haplogroup H and J cybrids. However, haplogroup H cybrids exhibited a degree of resistance to ATP depletion induced by tolcapone; a resistance that was not observed in haplogroup J cybrids. This agrees with findings reported by Ghelli *et al*., 2009 in which a cybrid model (non-hepatic) was used to determine that haplogroup J (vs haplogroups H and U) was more susceptible to uncoupling by the neurotoxic metabolite, 2,5-hexanediol (Ghelli et al., 2009). When assessing the relevance of the results from the present study, it is important to note that test concentrations were selected in order to generate the maximum effect on mitochondrial function in the absence of toxicity. The authors recognise that higher concentrations were used than C_max_ of compounds, however the goal was to model the mitochondrial toxicity experienced in a small number of individuals, which was not possible at lower concentrations. It is important to note that there was a lack of these described haplogroup-specific effects when using the non-hepatotoxic counterparts of the test compounds, bicalutamide and entacapone, illustrating that any differences are mechanisms- via induced mitochondria dysfunction.

Further analysis of the effects of the compounds upon respiration showed that the impact of flutamide and 2-hydroxyflutamide on parameters of mitochondrial respiration was similar between haplogroup H and J, though haplogroup J cybrids had consistently higher rates of respiration. Parameters of mitochondrial respiration also remained similar between the two cybrid haplogroups upon treatment with tolcapone. Given that the ATP level of cells, and indeed overall respiration, is the product of a myriad of processes to maintain energy status, mitochondrial function was dissected further by assessing the basal activity of specific respiratory complexes. Despite marginal differences in overall basal mitochondrial respiration, the basal activity of specific respiratory complexes showed significantly higher complex I and II activity in haplogroup J compared with haplogroup H cybrids. However, haplogroup J cybrids also displayed a heightened susceptibility to inhibition of complex I and II activity by flutamide/2-hydroxyflutamide. In contrast to complex I, complex II is entirely encoded in the nuclear genome, therefore one might predict that this should not differ between mitochondrial haplogroups. Nonetheless, upon the introduction of the foreign mitochondrial genome into cybrids, the initiation of retrograde responses to the nucleus must ensue, primarily via calcium signalling, which would enable regulation of the nuclear-encoded mitochondrial proteome and potentially enhanced biogenesis of respiratory complexes in haplogroup J (Luo et al., 1997; Amuthan et al., 2002; Srinivasan et al., 2015). This mechanism could account for the observed differential complex II activity.

The finding that haplogroup H cybrids are more resistant to ATP depletion induced by tolcapone offers new insights into the potential mechanisms underlying the onset of iDILI in certain individuals. In the case of tolcapone, four instances of liver failure among the 100 000 individuals who were administered the drug led to a black box warning and a switch in use to an alternative in-class compound, entacapone. However, as entacapone has been reported to be less efficacious than tolcapone, the ability to stratify therapy based upon the risk of iDILI, would be of great value (Rivest et al., 1999; Olanow, 2000; Watkins, 2000; Benabou and Waters, 2003; Olanow and Watkins, 2007; Lees, 2008; Longo et al., 2016).

Overall, this research offers insights into the potential importance of mitochondrial haplogroup on drug-induced mitochondrial dysfunction and iDILI. In the present study, HepG2 cybrids have been generated from only two mitochondrial haplogroups due to sample size limitations, but the method established offers itself to the generation of HepG2 cybrids from volunteers of other mitochondrial haplogroups and also from volunteers who are identical at the level of subhaplogroup. Furthermore, it is of note that differences in susceptibility have been observed despite the division of cybrids into two haplogroups which encompass much variation, and suggests that the comparison of more homogenous mtDNA subhaplogroups may provide further, more valuable insights into the differences conferred by differential mitochondrial background.

In conclusion, this study has established a novel, *in vitro* model that provides a preclinical representation of interindividual variation underpinning iDILI, thereby offering much greater translatability to clinical scenarios compared with current, homogenous preclinical models. This is paramount in understanding the safety of a drug in a range of populations prior to clinical trials, thus improving patient safety, as well as reducing drug attrition.

## Materials and Methods

### Materials

All forms of DMEM were purchased from Life Technologies (Paisley, UK). HepG2 cells were purchased from European Collection of Cell Cultures (Salisbury, UK). Cytotoxicity detection kits were purchased from Roche Diagnostics Ltd (West Sussex, UK). Clear and white 96-well plates were purchased from Fisher Scientific (Loughborough, UK) and Greiner Bio-One (Stonehouse, UK) respectively. All XF assay consumables were purchased from Agilent Technologies (CA, USA). All other reagents and chemicals were purchased from Sigma Aldrich (Dorset, UK) unless otherwise stated.

### Cohort

Ten healthy volunteers were recalled to give blood from the previously established HLA-typed archive (Alfirevic et al. 2012). These volunteers were selected upon the basis of their mitochondrial haplogroup (haplogroup H and J) and this genotyping has been described in our previous publication (Ball *et al*, 2021). Mitochondrial subhaplogroups and SNPs of the DNA isolated from each individual are described in the Supplementary Information. Ten donors were selected as an adequate number for a proof of principle study on the feasibility of generating the transmitochondrial cybrids. The volunteers were eligible to take part in the study if they were aged between 18 and 60 years, healthy and willing to donate one or more blood samples. The following exclusion criteria were applied and volunteers were not recruited if: they donated blood to transfusion services in the last 4 months; they had any medical problems, including asthma, diabetes, epilepsy or anemia; on any medications or if they had taken any recreational drugs in the last 6 weeks (including cannabis, speed, ecstasy, cocaine, LSD, and so on). Women were excluded if pregnant. This project was approved by the North West of England Research Ethics Committee and all participants gave written informed consent. Volunteer confidentiality was maintained by double coding DNA samples and by restricting access to participant’s personal data to trained clinical personnel. Detailed study eligibility and exclusion criteria have been published previously (Alfirevic et al. 2012).

### Generation of HepG2 cybrids

#### Generation of HepG2 p0 cells

HepG2 cells (ECACC Cat# 85011430, RRID:CVCL_0027) (≤ passage 7) were cultured and passaged as required in HepG2 ρ0 cell medium (DMEM/F-12 +GlutaMAX™ supplemented with FBS [10% v/v], L-glutamine [4 mM], sodium pyruvate [1 mM], HEPES [2 mM] and uridine [500 μM]) in the presence of ethidium bromide (EtBr; 1 μM). During treatment with EtBr (1 μM), the chemical was removed for 48 h every two weeks to help maintain cell viability. Following eight weeks’ exposure, EtBr was removed from a subset of cells for one week prior to characterisation to ensure cells were devoid of mtDNA i.e. were ρ0 cells (see Supplementary Information). If a ρ0 cell phenotype was not evident, cells were returned to EtBr treatment (1 μM) prior to removal for another week and re-testing, until a ρ0 cell population was observed.

#### Platelet isolation

Healthy volunteers, five of mitochondrial haplogroup H, and five of haplogroup J, donated whole blood from which platelets were isolated according to methods previously described (Ball et al., 2021). Briefly, 50 mL of blood was donated by each volunteer, and this fresh whole blood was immediately processed by a series of density centrifugation steps to produce isolated platelets. Throughout the procedure, PGI_2_ was used (1 μg/mL) to prevent platelet activation.

#### Platelet fusion with HepG2 rho zero (ρ0) cells

During the final centrifugation step of platelet isolation, HepG2 ρ0 cells were collected by trypsinisation and resuspended in HepG2 ρ0 cell medium. Following cell viability assessment (trypan blue; all viabilities were recorded at >90%), ρ0 cells (6 x 10^6^ cells) were centrifuged (1000 g, 5 min) and resuspended in Ca^2+^-free DMEM (2 mL; supplemented with 1 mM PGI2). This cell suspension was added to isolated platelets using a Pasteur pipette so as to minimise disruption to the platelet pellet. The cell mixture was then centrifuged (180 g, no brake, 10 min) to form a multi-layered pellet of platelets and HepG2 ρ0 cells.

Following centrifugation, the supernatant was removed and polyethylene glycol (250 μL; PEG 50%) was added before resuspending the cell pellet (30 s) followed by a 1 min incubation period. At the end of the incubation period, HepG2 ρ0 cell medium (30 mL) was added and a further 10-fold or 2-fold dilution with HepG2 ρ0 cell medium was performed before seeding into 96-well, 12-well and 6-well plates (Figure 7)

**Figure 7.**
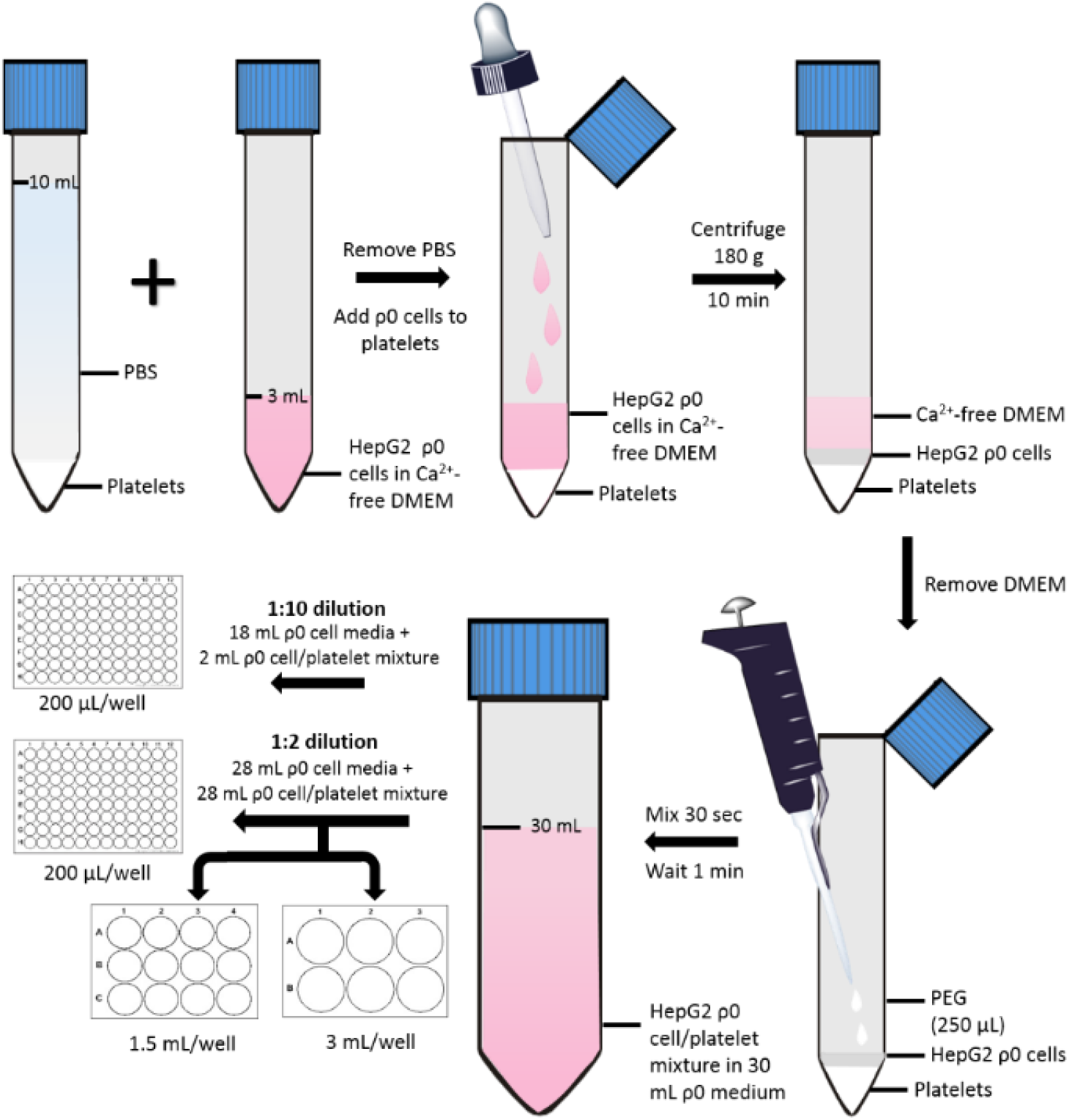
Schematic representation of the fusion of HepG2 ρ0 cells and platelets to generate HepG2 transmitochondrial cybrids. HepG2 ρ0 cells were assessed for viability (trypan blue; all viabilities were recorded at >90%), 6 x 10^6^ ρ0 cells were then centrifuged (1000 g, 5 min) and resuspended in Ca^2+^-free DMEM (2 mL; supplemented with 1 mM PGI_2_). This cell suspension was added to isolated platelets using a Pasteur pipette so as to minimise disruption to the platelet pellet. The cell mixture was then centrifuged (180 g, no brake, 10 min) to form a multi-layered pellet of platelets and HepG2 ρ0 cells. Following centrifugation, the supernatant was removed and polyethylene glycol (250 μL; PEG 50%) was added before resuspending the cell pellet (30 sec) followed by a 1 min incubation period. At the end of the incubation period, HepG2 ρ0 cell medium (30 mL) was added and a further 10-fold or 2-fold dilution with HepG2 ρ0 cell medium was performed before seeding into 96-well, 12-well and 6-well plates. Abbreviations: PEG, polyethylene glycol; ρ0, rho zero.

Two days post-fusion, media were replaced with fresh HepG2 ρ0 cell medium. After a further two days, this was replaced with medium consisting of equal volumes of HepG2 ρ0 medium and cybrid selection medium (high-glucose DMEM [glucose; 25 mM] supplemented with dialysed FBS [10% v/v], amphotericin B [1.35 μM] and antibiotic/antimycotic solution [100 units penicillin/mL, 170 μM streptomycin and 270 nM amphotericin B]). Finally, two days later, media were switched to 100% cybrid selection medium. ρ0 cells are auxotrophic for pyruvate and uridine, so the absence of these two constituents was the basis for the selection of successfully fused cells i.e. HepG2 cybrids. Cells remaining after selection were characterised to ensure a HepG2 cybrid phenotype by measuring the expression and function of mtDNA-encoded proteins (see Supplementary Information). Cells were then cultured and passaged as required in cybrid maintenance medium (DMEM high-glucose supplemented with FBS [10% v/v], L-glutamine [4 mM], sodium pyruvate [1 mM] and HEPES [1 mM]). For a schematic representation of HepG2 cybrid generation see Figure 8.

**Figure 8.**
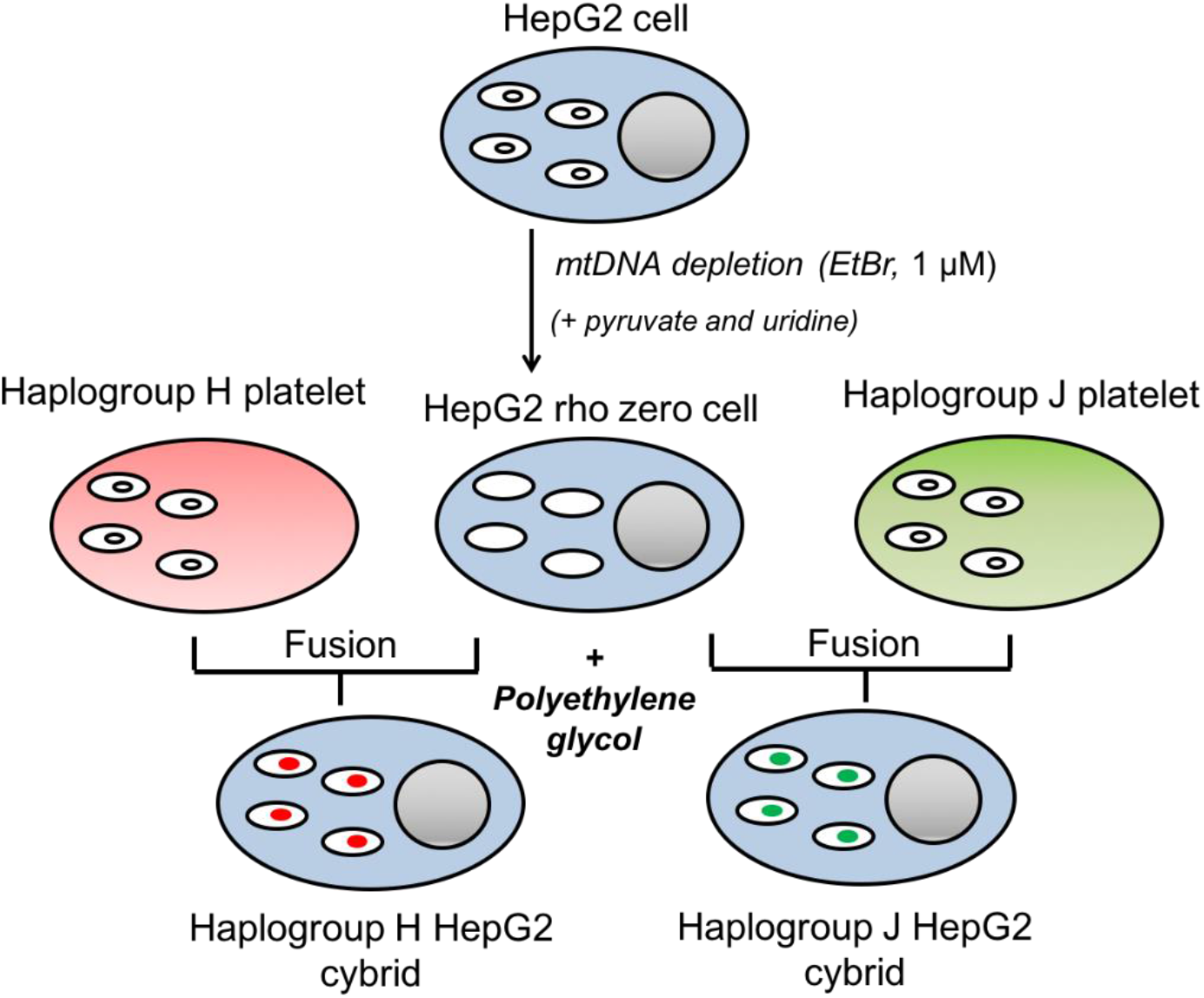
Schematic representation of HepG2 transmitochondrial cybrid generation. HepG2 cells were cultured in the presence of 1 μM ethidium bromide to generate HepG2 ρ0 cells. HepG2 ρ0 cells were combined with freshly-isolated platelets from healthy volunteers of mitochondrial haplogroup H or J and centrifuged (180 g, 10 min) to generate a multi-layered pellet. Following removal of the supernatant, polyethylene glycol (PEG; fusion reagent) was added and suspended with the cell mixture. After incubation with PEG for 1 min, 30 mL of HepG2 ρ0 cell medium was added (containing uridine and pyruvate) before further dilutions into a range of cell culture vessel sizes. This cell mixture was then cultured in cybrid selection medium (devoid of pyruvate and uridine), remaining cells were characterised to ensure the expression and function of mtDNA-encoded proteins. Abbreviations: EtBr, ethidium bromide; mtDNA, mitochondrial DNA.

### Assessment of mitochondrial function at basal state and following incubation with flutamide, 2-hydroxyflutamide and tolcapone

#### Dual assessment of mitochondrial function (ATP content) alongside cytotoxicity (LDH release)

##### Cell and reagent preparation

HepG2 cybrids were collected by trypsinisation and seeded on a collagen-coated flat-bottomed 96-well plate in cybrid maintenance medium (20 000 cells/50 μL/well) and incubated (24 h, 37°C, 5% CO_2_). Cells were then washed three times in serum-free galactose medium (DMEM containing 10 mM galactose and 6 mM L-glutamine) before addition of galactose medium (50 μL) and further incubation (2 h, 37°C, 5% CO_2_). This acute metabolic modification has been shown to be sufficient to allow the identification of drugs which induce mitochondrial dysfunction, by reducing the ATP yield from glycolysis, thereby increasing reliance on OXPHOS for ATP production. Flutamide, 2-hydroxyflutamide, tolcapone, bicalutamide and entacapone were each serially diluted to generate a concentration range of 0.01 – 300 μM in galactose medium. Diluted compounds (50 μL) were then added to each well (total well volume; 100 μL) and cells were incubated (2 h, 37°C, 5% CO_2_) before conducting assays to assess mitochondrial function and cytotoxicity. All assays used ≤0.5% DMSO as a vehicle control.

##### ATP content assay

ATP content was assessed by the addition of cell lysate (10 μL) and ATP standard curve solutions to a white-walled 96-well plate. ATP assay mix (40 μL; prepared according to the manufacturer’s instructions) was then added and bioluminescence was measured (Varioskan, Thermo Scientific).

##### LDH release assay

LDH release was determined by the extraction of 25 μL supernatant and 10 μL cell lysate from each well, before use of a cytotoxicity detection kit and reading at 490 nm. LDH release was calculated as: LDH supernatant/ (supernatant + lysate).

##### Normalisation (BCA assay)

Protein content was determined using cell lysate (10 μL) and protein standards (10 μL). BCA assay fluorescence was then measured at 570 nm.

#### Extracellular flux analysis

HepG2 cybrids were collected by trypsinisation and seeded on a collagen-coated XFe96 cell culture microplate (25 000 cells/100 μL medium/well; 96-well plate) and incubated (37°C, 5% CO_2_) overnight.

##### Mitochondrial stress test

Please refer to Ball et al, 2016 for a detailed description of this method (Ball et al., 2016). Briefly, cells were incubated (1 h, 37°C, 0% CO_2_) before replacement of culture medium with 175 μL of unbuffered Seahorse XF base medium supplemented with glucose (25 mM), L-glutamine (2 mM), sodium pyruvate (1 mM), pre-warmed to 37°C (pH 7.4). Following an equilibration period, measurements were taken to establish a baseline oxygen consumption rate (OCR) prior to the acute injection of each of the five test compounds (7.8-500 μM). Following compound injection a mitochondrial stress test consisting, of sequential injections of oligomycin (ATP synthase inhibitor; 1 μM), carbonyl cyanide 4-(trifluoromethoxy) phenylhydrazone (FCCP) (uncoupler; 0.5 μM) and rotenone/antimycin A (complex I/III inhibitors respectively; 1 μM each), was performed.

##### Respiratory complex assays

Please refer to Salabei et al, 2014 for a detailed description of this method (Salabei et al., 2014). Briefly, culture medium was replaced with mitochondrial assay solution buffer (MAS: MgCl_2_; 5 mM, mannitol; 220 mM, sucrose; 70 mM, KH_2_PO_4_; 10 mM, HEPES; 2 mM, EGTA; 1 mM, BSA; 0.4% w/v) containing constituents to permeabilise cells and stimulate oxygen consumption via complex I (ADP; 4.6 mM, malic acid; 30 mM, glutamic acid; 22 mM, BSA; 30 μM, PMP; 1 nM), complex II (ADP; 4.6 mM, succinic acid; 20 mM, rotenone; 1 μM, BSA; 30 μM, PMP; 1 nM), complex III (ADP; 4.6 mM, duroquinol; 500 μM, rotenone; 1 μM, malonic acid; 40 μM, BSA; 0.2% w/v, PMP; 1 nM) or complex IV (ADP; 4.6 mM, ascorbic acid; 20 mM, TMPD (N,N,N’,N’-tetramethyl-p-phenylenediamine); 0.5 mM, antimycin A; 2 mM, BSA; 30 mM, PMP; 1 nM). Following a basal measurement (no equilibration period) of three cycles of mix (30 s), wait (30 s) and measure (2 min), flutamide or 2-hydroxyflutamide (or MAS buffer for determination of basal complex activity) was injected followed by a mitochondrial stress test as detailed previously, only each measurement cycle was 3 min rather than 6 min.

#### Statistical analysis

In total 10 distinct cybrid cell lines were generated, one from each recruited volunteer, specifically 5 x distinct haplogroup H cell lines, and 5 x distinct haplogroup J cell lines. Additionally, 5 cybrid populations were generated for each cell line (volunteer). Each population was tested as an independent experiment (n =1), therefore giving a total of n=5 for each cybrid cell line/volunteer. Each independent experiment contained a minimum of 3 technical replicates.

Platelets were provided to the investigator blinded to haplogroup, to avoid bias; therefore, cybrid generation, experiments and subsequent data analyses were performed on cybrids for which the haplogroup was unknown. Unblinding occurred at the stage at which datasets were combined to the to enable the subsequent statistical comparison of haplogroup H vs haplogroup J.

Parameters for comparison were predefined to discourage statistical bias during analyses. These were, for both the assessment of mitochondrial function and drug-induced mitochondrial dysfunction: basal, maximum and ATP-linked respiration, spare respiratory capacity, proton leak and complex I-IV activity. Normality was assessed using a Shapiro-Wilk test. All data were assessed as parametric and therefore statistical significance was determined by an unpaired t-test using GraphPad Prism Version 7.0. Significance was determined when p value <0.05. EC50 data were determined by nonlinear regression analysis using GraphPad Prism 7.0 for the assessment of drug treatment on ATP level.

## Acknowledgements

The authors would like to thank the Royal Liverpool Research Facility, in particular, Lisa Gaskell, for the recruitment of volunteers and sample collection, and Prof. Dr. Peter Seibel and colleagues for their assistance in the generation of rho zero cells.

## Authors’ contributions

Author A.L.B generated the cybrid models, contributed to study design, completed laboratory work and data analysis, and wrote the manuscript. Author C.E.J contributed towards the generation of cybrid models and study design, and provided critical feedback. Author J.J contributed towards project conceptualisation. Author A.A contributed to funding acquisition, project conceptualisation and securement of resources including collaboration with the Royal Liverpool Research Facility. Author A.E.C acquired funding, conceptualised the project, directed data analysis and provided critical feedback. All authors have approved the final manuscript.

## Ethics approval statement

All procedures performed in studies involving human participants were in accordance with the ethical standards of the North West of England Research Ethics Committee (Cell Archive of HLA Typed Healthy Volunteers (HLA), CRN ID 7787, IRAS ID: 15623) with the 1964 Helsinki declaration and its later amendments or comparable ethical standards.

## Data availability

Source data are provided as files linked to the appropriate table/figures.

## Supplementary Information

### Mitochondrial DNA (MtDNA) variation of 10 healthy volunteers whose platelets were used to generate cybrids

| Donor | Haplogroup | Subhaplogroup | SNPs characteristic of assigned haplogroup | Additional SNPs |
| --- | --- | --- | --- | --- |
| 1 | H | H1c3 | 257G 263G 477C 750G 1438G 3010A<br>4769G 8473C 8860G 15326G | 195C 12966T<br>16519C |
| 2 | H | H1bb | 152C 263G 750G 1438G 3010A 4769G<br>8860G 11864C 15326G | 16519C |
| 3 | H | H1a1 | 73G 263G 750G 1438G 3010A 4769G<br>6365C 8860G 15326G 16162G | 152C 3483A<br>16360T<br>16519C |
| 4 | H | H1c | 263G 477C 750G 1438G 3010A 4769G<br>8860G 15326G | 10646A<br>16519C |
| 5 | H | H2a1e1a1 | 263G 575T 750G 751G 951A 8860G<br>15326G 16124C 16148T 16166G 16354T |  |
| 6 | J | J2b1g | 73G 150T 152C 263G 489C 750G 1438G<br>4769G 5633T 7028T 8860G 9872G<br>10172A 10398G 11251G 11719A 12612G<br>13708A 14766T 15257A 15326G 15452A<br>15812A 16069T 16126C 16193T | 2789T 13821T |
| 7 | J | J1c+16261 | 73G 185A 228A 263G 295T 462T 489C<br>750G 1438G 3010A 4216C 4769G 7028T<br>8860G 10398G 11251G 11719A 12612G<br>13708A 14766T 14798C 15326G 15452A<br>16069T 16126C 16261T | 4113A 6554T<br>10915C<br>16316G |
| 8 | J | J1c1c | 185A 228A 462T 482C 489C 750G 1438G<br>3010A 3394C 4216C 4769G 7028T<br>8860G 10398G 11251G 11719A 13708A<br>14766T 14798C 15326G 15452A 16069T<br>16126C 16145A | 188G 13943T<br>14552G |
| 9 | J | J1c2h | 185A 188G 222T 228A 263G 295T 462T<br>489C 750G 1438G 3010A 4216C 4769G<br>7028T 8860G 10398G 11251G 11719A<br>12612G 13708A 14766T 14798C 15326G<br>15452A 16069T 16126C | 16189C<br>16519C<br>16527T |
| 10 | J | J1c1e | 185A 228A 263G 295T 462T 482C 489C<br>750G 1438G 3010A 3394C 4216C 4769G<br>7028T 8860G 10398G 10454C 11251G<br>11719A 12612G 13708A 14766T 14798C<br>15326G 15452A 16069T 16126C 16368C | 13889A |

Additional SNPs refers to SNPs that were present in the sample but were not characteristic of the assigned haplogroup. Abbreviations: SNP, single nucleotide polymorphism.

### Characterisation of HepG2 rho zero cells and HepG2 transmitochondrial cybrids

#### S1 Cell doubling time

The dependence of HepG2 ρ0 cells on pyruvate and uridine meant that culture media devoid of these constituents was able to select for the successful depletion of mtDNA and the resultant non-functional electron transport chain in EtBr-treated cells. Concordantly, HepG2 WT cells had a similar doubling time in media with or without these additives, averaging 26.2 hours. In contrast, HepG2 ρ0 cells exhibited no growth without uridine and pyruvate and an average doubling time of 27.6 hours

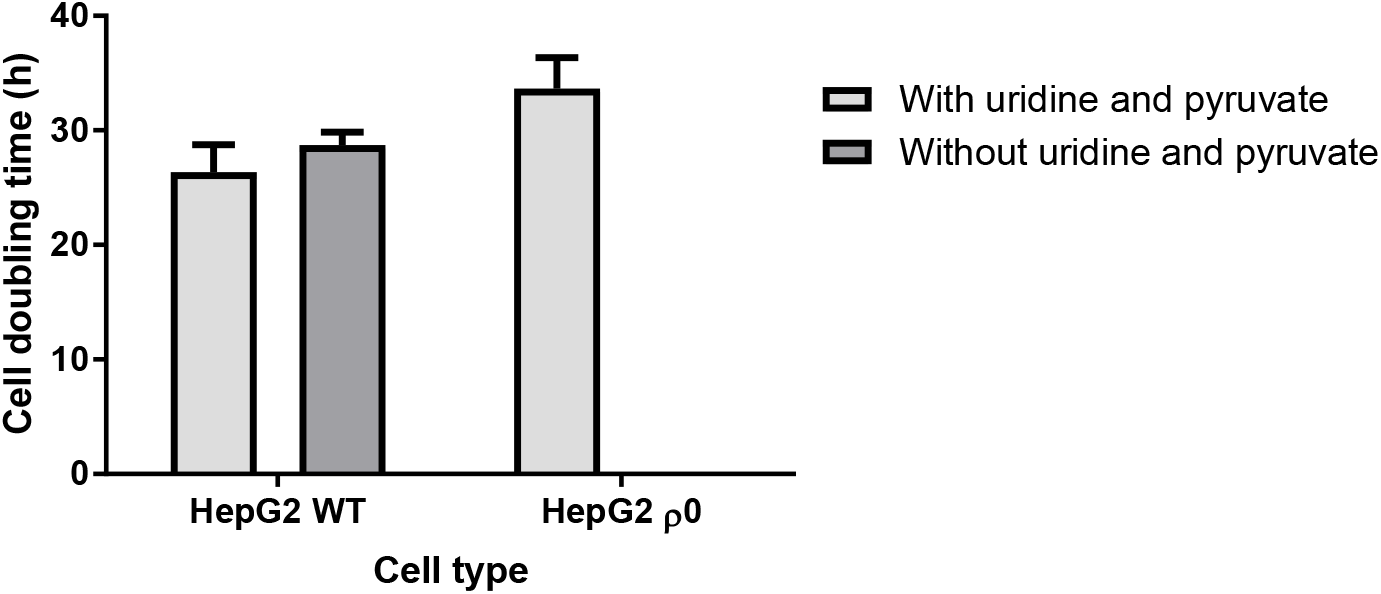

when in media containing these two constituents.

**Cell doubling time of HepG2 wild-type (WT) and HepG2 rho zero (ρ0) cells.** The two cell types were cultured in media with or without uridine and pyruvate and growth rate calculated. Data are presented as mean+SEM of n=3 experiments. Source data: figs s1 – source data file.xslx

#### S2 Mitochondrial DNA content

The Ct value of both mtDNA primers increased dramatically in ρ0 compared to WT cells whilst nuclear DNA Ct values remained consistent across all three cell lines. Similarly, both HepG2 WT and cybrid cells had thousands of mtDNA copies/cell in contrast to the ρ0 cells which had less than one copy/cell

Assessment of mtDNA content in HepG2 wild-type (WT), HepG2 rho zero (ρ0) and HepG2 cybrids.

**Assessment of mtDNA content in HepG2 wild-type (WT), HepG2 rho zero (p0) and HepG2 cybrids.**
| Sample | Primer C<sub>t</sub> value |  |  |  | mtDNA copies/cell |  |  |  |
| --- | --- | --- | --- | --- | --- | --- | --- | --- |
|  | TERT | RNase P | ND-1 | Custom | TERT/ND-1 | TERT/custom | RNase P/ND-1 | RNase P/custom |
| HepG2 WT | 30.6<br>(1.00) | 31.3<br>(1.11) | 18.5<br>(0.120) | 18.9<br>(0.140) | 9280 | 6985 | 14563 | 10960 |
| HepG2 p0 | 27.7<br>(0.0600) | 27.2<br>(0.0700) | 34.7<br>(0.740) | 32.0<br>(0.180) | 0.00400 | 0.0240 | 0.00300 | 0.0170 |
| HepG2 | 29.8<br>(1.10) | 30.4<br>(0.550) | 18.9<br>(0.730) | 19.0<br>(0.0900) | 3875 | 3590 | 5673 | 5256 |

| cybrid |
| --- |

Abbreviations: C<sub>t</sub>, cycle threshold; ND-1, NADH dehydrogenase-1; RNase, ribonuclease P RNA component H1; TERT, telomerase reverse transcriptase; WT, wild-type; p0, rho zero. C<sub>t</sub> values are displayed as mean (SEM).

#### S3 Detection of mitochondrial/nuclear DNA-encoded mitochondrial proteins

Western blot analysis showed expression of all nuclear DNA and mtDNA-encoded subunits of the electron transport chain which were probed for in HepG2 WT and cybrid cells. However, ρ0 cells did not express the mtDNA-encoded subunit of complex IV. Notably, despite all the other subunits being encoded in the nuclear DNA, it was only the alpha subunit of ATP synthase that was retained in the ρ0 cells.

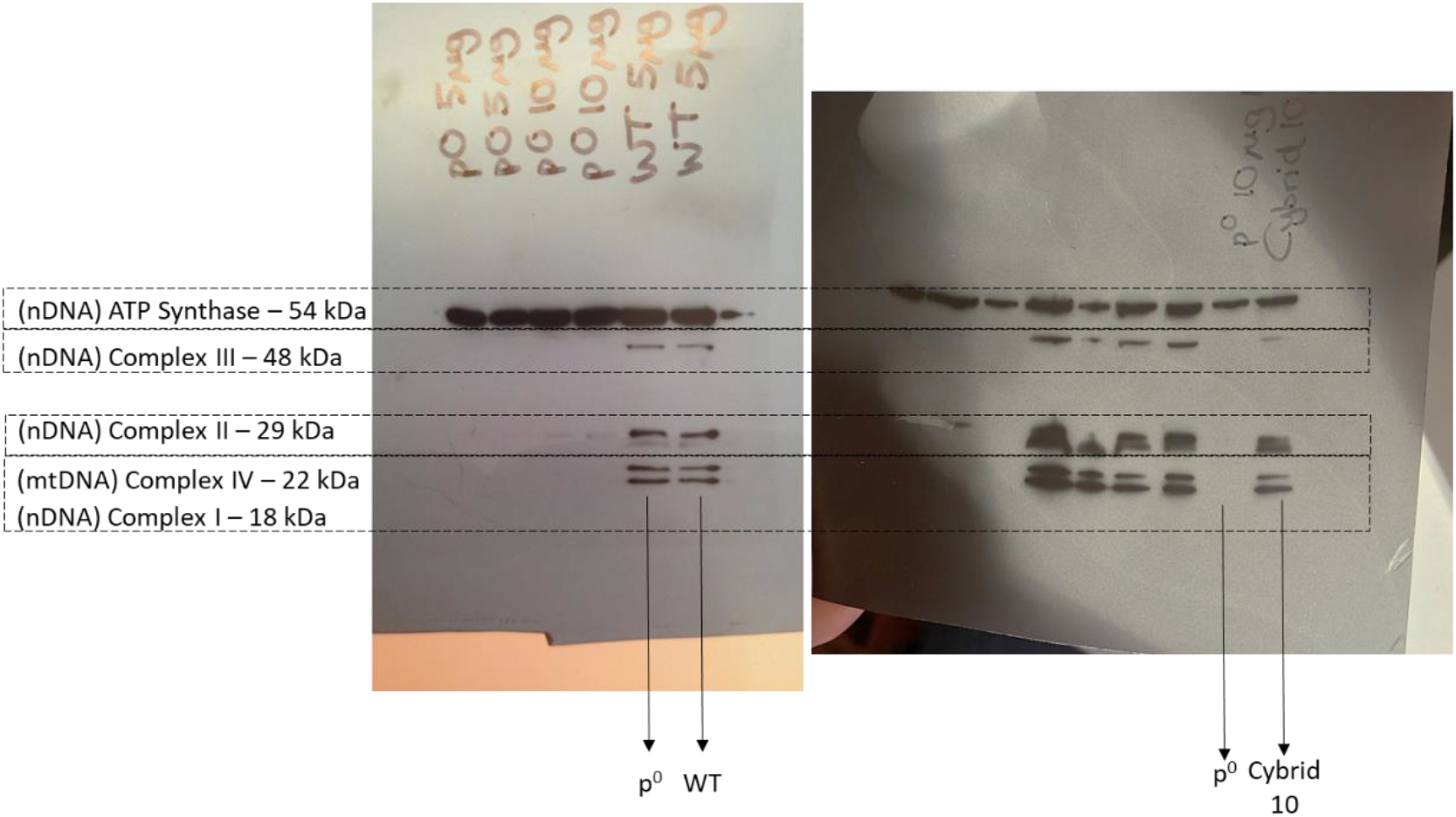

**Representative western blots of HepG2 wild-type (WT), rho zero and cybrid cell lysates.** 10 μg of lysate protein was resolved by SDS-PAGE and probed for subunits of complexes I (NDUFB8), II (Iron-sulphur protein (IP) 30 KDa), III (Core 2), IV (II), V (alpha). Abbreviations: mtDNA, mitochondrial DNA; nDNA, nuclear DNA. Source data: figs s3 and s4 source data file.xslx and figs s3 –raw data.pptx

#### S4 Detection of electron transport chain function

HepG2 WT and cybrid cells exhibited a classic response to the series of mitochondrial inhibitors used to perform the mitochondrial stress test whereas the ρ0 cells did not respond to these inhibitors and had very low basal OCR, all of which was due to non-mitochondrial respiration. The PPRgly /OCR ratio was also higher in WT and ρ0 cells compared with cybrid cells.

Differences in parameters of mitochondrial function in HepG2 wild-type (WT), rho zero (ρ0) and cybrid cells.

**Differences in parameters of mitochondrial function in HepG2 wild-type (WT), rho zero (p0) and cybrid cells.**
| HepG2 cell type | Basal OCR (pmol/min/µg protein) | PPR<sub>gly</sub>/OCR | % Non-mitochondrial respiration |
| --- | --- | --- | --- |
| WT | 6.52 (0.350) | 0.354 (0.0900) | 28.4 (0.310) |
| Rho zero | 0.450 (0.100) | 55.5 (0.930) | 102 (4.33) |
| Cybrid | 6.53 (0.400) | 0.466 (0.660) | 24.3 (0.510) |

Abbreviations: OCR, oxygen consumption rate; PPR<sub>gly</sub>, proton production rate attributed to glycolysis; WT, wild-type.
Values are displayed as mean (SEM). Source data: figs s3 and s4 source data file.xlsx

#### S5 The effect of bicalutamide upon haplogroup H and J transmitochondrial cybrids

Bicalutamide-treated cybrids of each haplogroup exhibited a similar decline in ATP content until the highest concentration used (300 μM), when haplogroup J cybrids had significantly less ATP (Figure A). As with flutamide and 2-hydroxyflutamide treatment, the decline in ATP was in the absence of significant LDH release. No significant differences were evident between the two cybrids groups in parameters of mitochondrial function using XF analysis, however as was the case with flutamide and 2-hydroxyflutamide, haplogroup J exhibited higher proton leak in both control and treated cybrids (Figure B-F).

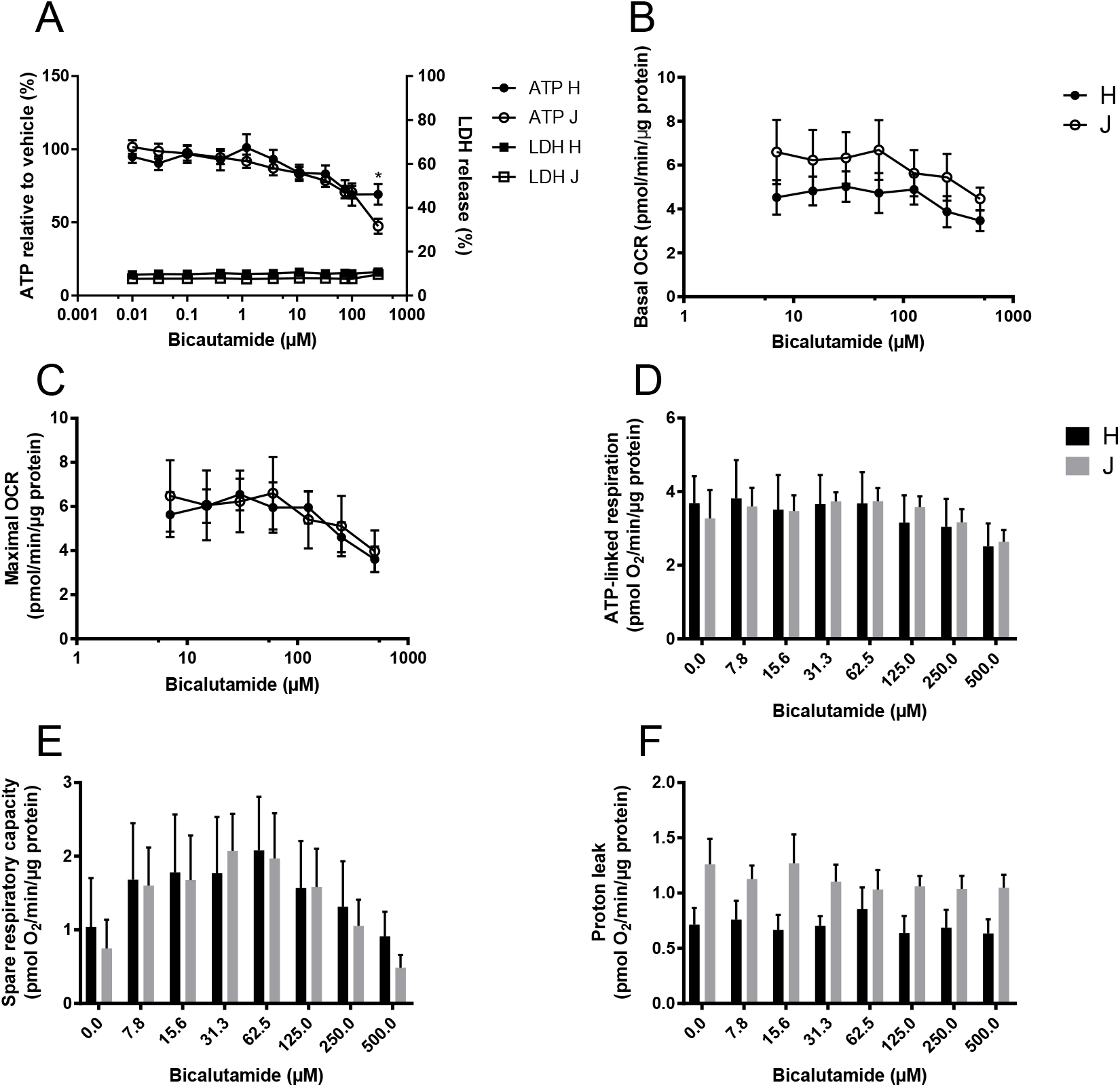

**The effect of bicalutamide upon ATP levels and mitochondrial respiratory function in haplogroup H and J HepG2 cybrids.**

**A:** Cybrids were treated (2 h) up to 300 μM bicalutamide in galactose medium. ATP values are expressed as a percentage of those of the vehicle control. Lactate dehydrogenase (LDH) release is expressed as extracellular LDH as a % of total LDH. **B-F:** XF analysis-detected changes in basal and maximal respiration, ATP-linked respiration, spare respiratory capacity and proton leak following acute treatment with bicalutamide (up to 500 μM). Statistical significance between haplogroup H and J cybrids: * *p*<0.05. Data are presented as mean ± SEM of n = 5 experiments. Source data: figs s5 – source data file.xslx

#### S6 The effect of entacapone upon haplogroup H and J transmitochondrial cybrids

Entacapone induced a weaker decline in ATP levels compared to tolcapone (Figure A). There was no significant difference in ATP levels between the two haplogroups and also no significant difference in parameters of mitochondrial function, though haplogroup H cybrids had consistently higher ATP-linked respiration and spare respiratory capacity which tended towards significance (Figure D, E).

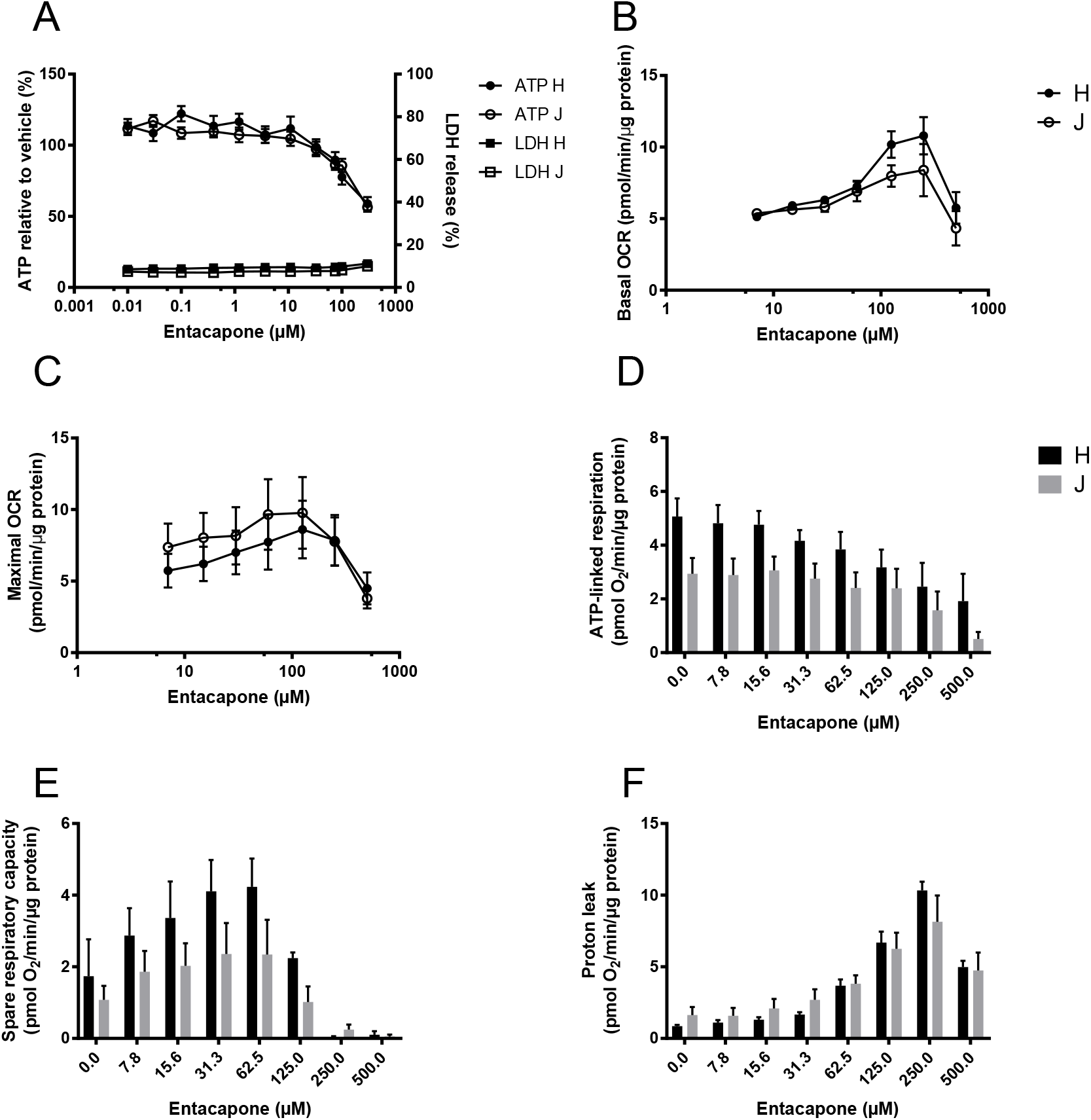

**The effect of entacapone upon ATP levels and mitochondrial respiratory function in haplogroup H and J HepG2 transmitochondrial cybrids. A:** Cybrids were treated (2 h) with serial concentrations up to 300 μM entacapone in galactose medium. ATP values are expressed as a percentage of those of the vehicle control. Lactate dehydrogenase (LDH) release is expressed as extracellular LDH as a % of total LDH. **B-F:** Changes in basal respiration, maximal respiration, ATP-linked respiration, spare respiratory capacity and proton leak following acute treatment with entacapone (up to 500 μM). Data are presented as mean ± SEM of n = 5 experiments. Source data: figs s6 – source data file.xslx

## Appendix Supplementary Methods

### Methodology

#### Cell doubling time

HepG2 wild-type (WT) cells and HepG2 rho zero (ρ0) cells were seeded at 30 000 cells/well in a 24-well plate in either ρ0 cell media (contains pyruvate and uridine) or selection media (devoid of pyruvate and uridine). On days 1, 3, 5 and 7 of culture, cells were collected by trypsinisation and counted, following which growth rate was calculated. Rho zero cells are auxotrophic for pyruvate and uridine, but WT cells are not, therefore the absence of cell growth in selection media indicated complete loss of mtDNA.

#### DNA extraction and real-time PCR

DNA extraction from HepG2 WT, HepG2 ρ0 and HepG2 cybrid cells was performed using a DNA mini kit (Qiagen, Manchester, UK) according to the manufacturer’s instructions. Sample DNA concentrations and quality were then quantified using a Quant-iT™ PicoGreen™ dsDNA Assay Kit and nanodrop spectrophotometry respectively (Fischer Scientific, Loughborough, UK).

Real-time PCR was carried out using two primers for regions of mtDNA; a custom sequence and ND-1 (complex I subunit) and two primers for regions of nuclear DNA; telomerase reverse transcriptase (TERT) and RNase P (Applied Biosystems, California, USA) (Malik et al., 2011).

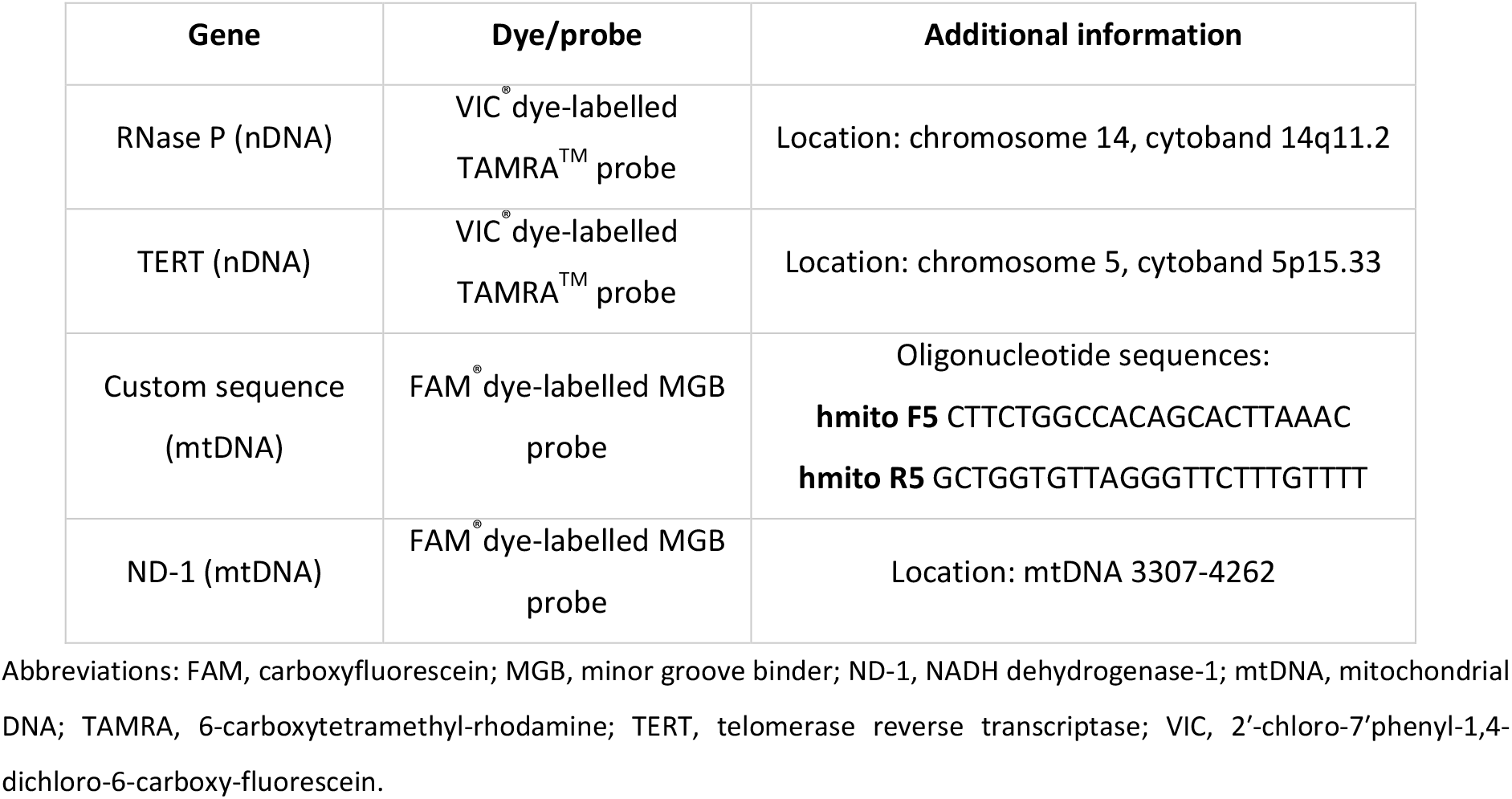
Real-time PCR primers used to amplify regions of mitochondrial and nuclear DNA.

During sample preparation, 2X Taqman^®^ genotyping master mix (5 μL), a nuclear DNA primer (0.5 μL), mtDNA primer (0.5 μL), dH_2_0 (2 μL) and 10 ng DNA (2 μL) from each sample were combined to give a final sample concentration of 1 ng/μL in each well. Real-time PCR was then carried out using the viiA7 RT-PCR system (Life Technologies, UK). MtDNA copies per cell were calculated on the basis that each nuclear DNA primer was present in diploid copies per cell and used the following formula, where x_1_= nuclear DNA primer cycle threshold (C_t_) value, x_2_=mtDNA primer C_t_ value: mtDNA copies per 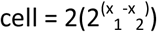 (Schäfer, 2016).

#### Detection of mitochondrial/nuclear DNA-encoded mitochondrial proteins

HepG2 ρ0, HepG2 WT or HepG2 cybrid cells were lysed using sonication and 10 μg of lysate protein was resolved by sodium dodecyl sulphate-polyacrylamide *gel* electrophoresis (SDS-PAGE) using 4-12% Bis-Tris gel (Invitrogen, UK) in MOPS buffer (MOPS; tris-base; 1.21% w/v, sodium dodecyl sulphate; 0.20% w/v, EDTA; 0.06% w/v in distilled water (dH_2_0)).

This gel was then transferred to a nitrocellulose membrane (GE Healthcare, Buckinghamshire, UK) in transfer buffer (tris-base; 0.30% w/v, glycine; 1.5% w/v, methanol; 20% v/v in dH_2_0) and blocked using 10% non-fat dried milk in Tris Buffered Saline-Tween (TBS-T: TBS; 0.50% v/v, tween; 0.10% v/v in dH_2_0).

Blocking solution was removed using TBS-T and the membrane probed for CI-20, CII-30, CIII-core2, CIV-I and CV-alpha subunits of complexes I-V of the electron transport chain using MitoProfile^®^ Total OXPHOS Human WB Antibody Cocktail (Abcam, Cambridgeshire, UK) (0.20% v/v in 10% non-fat dried milk in TBS-T). This was followed by anti-mouse secondary antibody (0.01% v/v in 10% non-fat dried milk in TBS-T) before visualisation using an ECL™ system (GE Healthcare, Buckinghamshire, UK).

#### Detection of electron transport chain function

Mitochondrial stress tests were performed on untreated HepG2 WT, p0 and HepG2 cybrid cells using extracellular flux analysis as described in the main text.

Extracellular flux analysis produced two raw outputs, oxygen consumption rate and extracellular acidification rate (ECAR). ECAR can be indicative of the glycolytic rate of the cell, however it also takes into account changes in ECAR due to oxidative phosphorylation. Therefore the measure of PPR_gly_, glycolytic production rate was used to quantify glycolysis, this was calculated by subtracting respiratory acidification contributions from the total proton production rate.

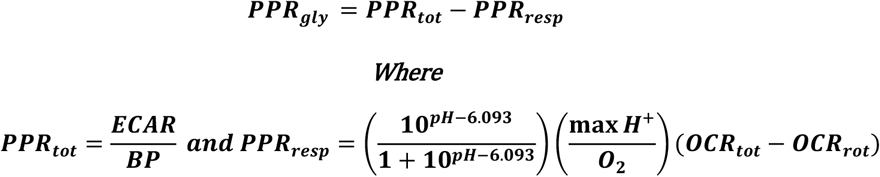

##### Equations for the calculation of PPR_gly_ from mitochondrial stress tests

Abbreviations: PPR_gly_, proton production rate attributed to glycolysis; PPR_resp_, proton production rate attributed to respiration; PPR_tot_, total proton production rate; ECAR, extracellular acidification rate; BP, buffering power; max H^+^/O_2_, derived acidification for metabolic transformation of glucose oxidation; OCR_tot_, total oxygen consumption rate; OCR_rot_, oxygen consumption rate following rotenone injection (Kelly, 2018).

